# Human hippocampal replay during rest prioritizes weakly-learned information and predicts memory performance

**DOI:** 10.1101/173021

**Authors:** Anna C. Schapiro, Elizabeth A. McDevitt, Timothy T. Rogers, Sara C. Mednick, Kenneth A. Norman

## Abstract

There is now extensive evidence that the hippocampus replays experiences during quiet rest periods, and that this replay benefits subsequent memory. A critical open question is how memories are prioritized for replay during these offline periods. We addressed this question in an experiment in which participants learned the features of 15 objects and then underwent fMRI scanning to track item-level replay in the hippocampus using pattern analysis during a rest period. Objects that were remembered less well were replayed more during the subsequent rest period, suggesting a prioritization process in which weaker memories—memories most vulnerable to forgetting—are selected for wake replay. Participants came back for a second session, either after a night of sleep or a day awake, and underwent another scanned rest period followed by a second memory test. In the second session, more hippocampal replay of a satellite during the rest period predicted better subsequent memory for that satellite. Only in the group with intervening sleep did rest replay predict improvement from the first to second session. Our results provide the first evidence that replay of individual memories occurs during rest in the human hippocampus and that this replay prioritizes weakly learned information, predicts subsequent memory performance, and relates to memory improvement across a delay with sleep.

## Introduction

The brain is highly active even when an organism is disengaged from its sensory environment [1]. There is accumulating evidence from the rodent literature that the hippocampus replays recent experiences during these rest periods, typically measured as place cells firing in a sequence corresponding to an experienced trajectory of locations [2, 3]. These replay events appear to be functional: they relate to later memory performance and their disruption impairs memory [4-6]. Many human studies, mainly using fMRI, likewise suggest that content from a recent experience reactivates during subsequent rest periods, and that this activity relates to later memory [7-17]. These studies have found replay^1^ of individual items outside the hippocampus [7-9] and category-level reinstatement within the hippocampus [10, 14], though replay of individual items within the human hippocampus has not yet been observed.

What process determines which memories get replayed? The brain likely cannot replay every experienced event during rest, nor would it be worthwhile to do so—not all memories need or deserve further processing. There is evidence that memories are more likely to be replayed during subsequent awake rest when associated with reward or fear [14, 15, 17]. There is also evidence that certain kinds of memories benefit more from a period of sleep, including memories that are relevant to future behavior [20], that are emotionally laden [21], and that are weaker in strength [22-29, cf. 30, 31]. Of particular relevance, this prioritization of weaker memories was observed in prior work using the current paradigm, where objects exposed least frequently during training benefitted the most from a nap [32]. These sleep studies do not directly demonstrate that replay prioritizes weaker (or other types of) memories, as replay was not measured, but the ubiquity of sleep replay and its association with memory improvement [18, 19] suggest that prioritized replay is a potential mediator of the behavioral benefits. While there have been many studies assessing the relationship of rest replay to subsequent memory, there has not yet been a direct test of how initial memory strength relates to subsequent replay.

The current study assesses how replay is prioritized on the dimension of memory strength, and tests the effects of such replay on subsequent memory, using a property-inference task developed in prior work [32]. Participants learned the features of 15 “satellite” objects belonging to three categories (Fig. 1), where satellites in the same category shared most of their features. Those assigned to a Sleep group participated in a first session in the evening and a second session the next morning, while those assigned to a Wake group had their first session in the morning and second session that evening. In Session 1, we (1) taught participants the features of the satellites, (2) tested their memory for these features, (3) measured the neural response generated by each of the satellites in the fMRI scanner, and then (4) used pattern analysis to assess whether individual items were replayed by the hippocampus during a rest period in the scanner. In the second session, we again tested memory, measured neural responses to the individual satellites, and assessed replay during rest, with the one difference that the memory test came last instead of first.

**Figure 1.**
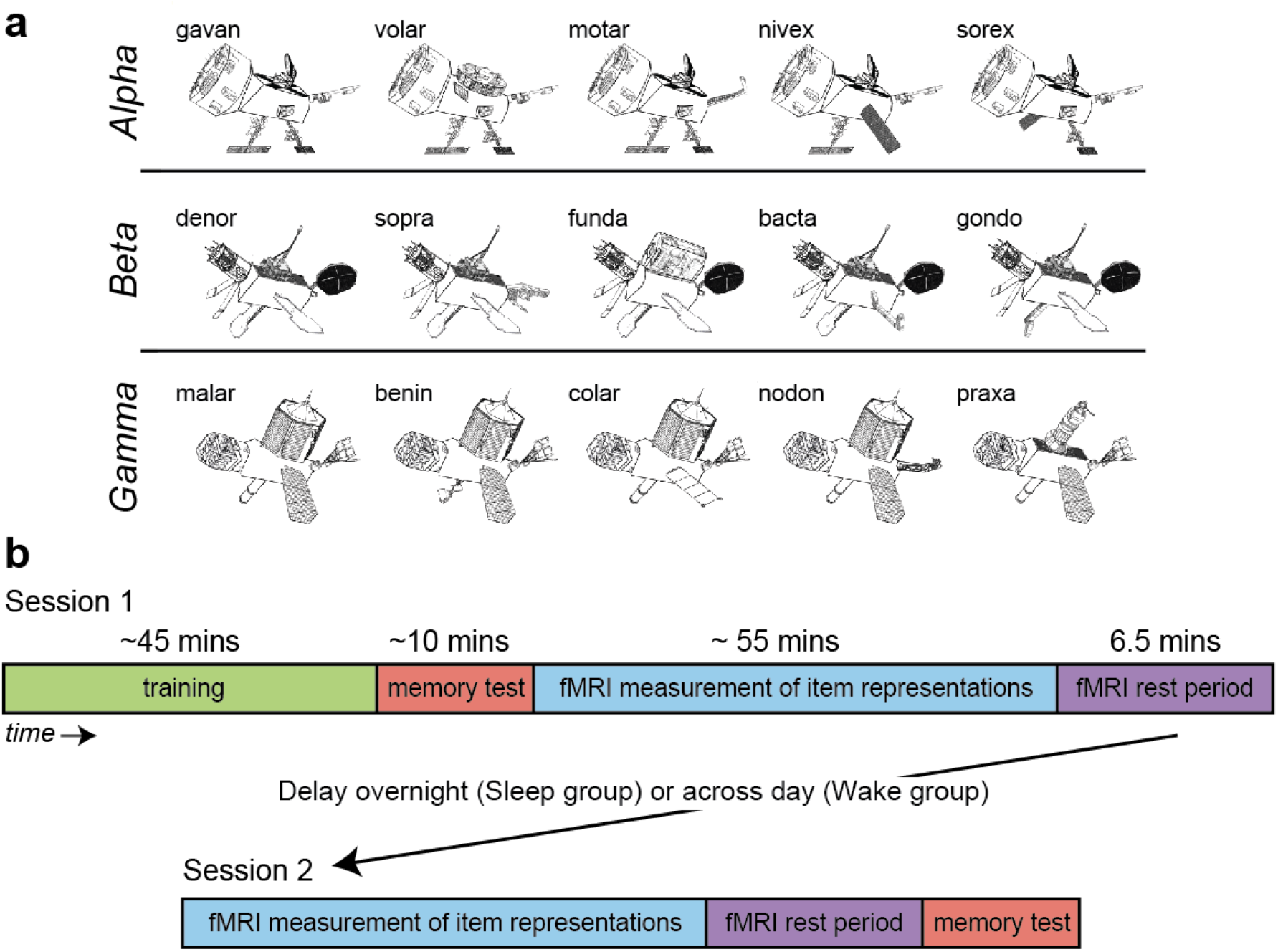
Stimuli and design. (**a**) Examples of stimuli presented from the three classes Alpha, Beta, and Gamma, labeled with unique code names. Satellites were built randomly for each participant, using the same category structure. (**b**) Session 1 and 2 procedures for all subjects, with delay between sessions either overnight or across day depending on group.

This design allowed us to answer four questions critical to understanding the role of hippocampus in the consolidation of object knowledge:

1. Are representations of recently learned individual items replayed in the human hippocampus during quiet rest? Prior literature in humans and rodents suggests that this occurs, but it has not yet been observed at the resolution of individual items in humans.

2. Does probability of replay relate to the strength of a memory, as determined by the initial memory test? It is possible that weaker memories are prioritized for replay, as suggested by the sleep literature; alternatively, stronger memories may be more likely to persist into subsequent rest.

3. Does replay of specific items in the hippocampus predict subsequent memory? Prior literature in humans and rodents suggests a positive relationship, though this has not yet been assessed for individual hippocampal memories in humans.

4. Does the relationship between wake replay and memory improvement over a delay relate to the presence of intervening sleep? Replay measured during awake rest periods may reflect (or perhaps influence [47, 48]) the processing that continues to occur in the intervening period between sessions, and this processing may be especially beneficial over sleep [33].

## Methods

### Participants

24 participants (14 females, mean age = 24.5 years, range = 19–38 years) from the Princeton University community participated in exchange for monetary compensation or course credit. Data from six additional subjects were excluded due to excessive motion (three subjects), technical issues (two subjects), and poor performance on the one-back task in the scanner (one subject; A' > 2 SD below average). Informed consent was obtained from all subjects, and the study protocol was approved by the Institutional Review Board for Human Subjects at Princeton University.

Subjects reported no history of neurological disorders, psychiatric disorders, major medical issues, or use of medication known to interfere with sleep. Subjects also reported having a regular sleep-wake schedule, which was defined as regularly going to bed no later than 2AM, waking up no later than 10AM, and getting at least 7 hours of total sleep per night on average. The Epworth Sleepiness Scale [ESS; 34], the Morningness-Eveningness Questionnaire [MEQ; 35], and the Pittsburgh Sleep Quality Index [PSQI; 36] were used to screen out potential subjects with excessive daytime sleepiness (ESS score >10), extreme chronotypes (MEQ < 31 or > 69), and poor sleep quality (PSQI > 5).

### Stimuli

Participants learned about 15 novel “satellite” objects organized into three classes (Fig. 1). Each satellite has a “class” name (Alpha, Beta, or Gamma) shared with other members of the same category, a unique “code” name, and five visual parts. One of the satellites in each category is the prototype (shown on the left for each category in Fig. 1), which contains all the prototypical parts for that category. Each of the other satellites has one part deviating from the prototype. Thus, each non-prototype shares 4 features with the prototype and 3 features with other non-prototypes from the same category. Exemplars from different categories do not share any features. Each satellite has *shared features*: the class name and the parts shared among members of the category, and *unique features*: the code name and the part unique to that satellite (except for the prototype, which has no unique parts). Satellites were constructed randomly for each participant, constrained by this category structure.

### Procedure: prior to experiment

Subjects were instructed to attempt at least 7 hours of sleep per night for 5 nights prior to (and during, in the case of Sleep subjects) the study. Adherence to the sleep schedule was tracked with daily sleep diaries. Subjects were also asked to refrain from any alcohol for 24 hours prior to the first session, and throughout the experiment, and to keep caffeine intake to a minimum during this period. Heavy caffeine users (>3 servings per day) were not enrolled.

### Procedure: Session 1 Training

Participants learned about the satellites in two phases. In the first phase, which lasted 15 minutes on average, the satellites were introduced one by one. For each satellite, the class and code name were displayed, followed by the image of the satellite. A box highlighted each of the five visual features on the satellite image one by one, to encourage participants to attend to each feature. Participants were then asked to recall the class and code names by clicking on one of three options given for each name. Next, participants used a point-and-click interface to try to reconstruct the satellite image from scratch. Icons representing the five part types were displayed on the right hand side of the screen, and when an icon was clicked, all the possible versions of that part were displayed in a row on the bottom of the screen. The participant could then click on one of the part versions on the bottom to add it to the satellite in the center of the screen. If the participant was too slow at this task (took longer than 15 s), or reconstructed the satellite incorrectly, a feedback screen would appear displaying the correct features.

In the second phase of training, which lasted 32 minutes on average, participants were shown a satellite with one feature missing, which could be one of the five visual features, the code name, or the class name (code and class name buttons were displayed along with the part icons on the right hand side of the screen, and when selected, displayed the corresponding name options in a row on the bottom). Using the same point-and-click interface, participants chose a feature (out of all possible) to complete the satellite. If they chose the correct feature, they were told it was correct, and could move on to the next trial. If they chose an incorrect feature, they were shown the correct feature, and had to repeat the trial until they chose the correct feature.

Remembering the shared properties of the satellites is easier than remembering the unique properties, as the shared properties are reinforced across study of all the satellites in the same class. The task was titrated in pilot testing to ensure that, at the end of training, participants performed equivalently at retrieving shared and unique properties of the satellites. To accomplish this, unique features were queried 24 times more frequently than shared features. This phase of training continued until the participant reached a criterion of 66% of trials correct on a block of 32 trials, or until 60 minutes had passed.

### Procedure: Session 1 Test

Immediately after training, participants were tested by again filling in missing features of the satellites, now without feedback. The test phase had 39 trials, with two missing features per trial, which allowed us to collect more information per trial as well as provide less exposure to the correct features (to minimize learning during the test phase). The test phase took 10 minutes on average. Each satellite appeared twice in the test phase: once with its code name and its class name or one shared part tested, and once with two shared parts or one unique part and one shared part tested. The remaining 9 trials tested generalization to novel satellites. Novel satellites were members of the trained categories but had one novel feature. The queried feature for novel items was always a shared part (class name or shared visual feature). Test trials were presented in a random order.

### Procedure: fMRI scanning

After completion of the first session test phase, participants were scanned while viewing the satellite images (without names) for 52 minutes. Satellites subtended up to 19 degrees of visual angle on the scanner projection. There were 8 runs, lasting 6.5 mins each, with self-paced breaks between runs. In each run, each of the 15 images was presented four times in pseudo-random order, such that each satellite appeared in the first, second, third, and fourth quarter of the trials. Four trials in each run were randomly chosen to be duplicated, such that these satellites were shown twice in immediate succession. These served as rare (4 trials out of 64) targets for a one-back task that subjects performed while viewing the satellites, to encourage maintenance of attention. Subjects pressed one key on a keypad to indicate that the current satellite was not an exact repetition of the previous satellite, and a different key to indicate that it was a repetition. Keys corresponded to index and middle fingers of the right hand, with key assignment for repetition and no-repetition counterbalanced across subjects. Feedback for responses at each trial was provided as a green or red dot at fixation. Each satellite was presented for 3s with a jittered interstimulus interval (40% 1s, 40% 3s, 20% 5s), to facilitate modeling of the response to individual items.

Next, we collected a ninth 6.5-min functional run where participants were instructed to relax and watch the fixation dot on the screen, emphasizing that they should keep their eyes open.

### Procedure: Session 2

In Session 2, participants did the same scan procedure, with images presented in a different random order. Then they got out of the scanner and completed the same test phase as in the first session, with items presented in a different random order. The Karolinska Sleepiness Scale [KSS; 37], which assesses state sleepiness/alertness on a scale of 1 (extremely alert) to 9 (very sleepy), was completed at the beginning and end of each session.

Participants in the Sleep group (n=12) began the first session around 7pm and the second session around 9am, and participants in the Wake group (n=12) began the first session around 9:30am and the second session around 10pm (time choices were constrained by scanner availability). They were not told in the first session that there would be a second memory test, though when asked afterwards, they generally reported feeling that a second test was likely and that they were not surprised by it. Subjects in the Wake condition were instructed not to nap between sessions.

### fMRI data acquisition

Data were acquired using a 3T Siemens Skyra scanner with a volume head coil. In each session, we collected 9 functional runs with a T2*-weighted gradient-echo EPI sequence (36 oblique axial slices: 3×3 mm inplane, 3 mm thickness; TE=30 ms; TR=2000 ms; FA=71°; matrix=64×64). Each run contained 195 volumes. We collected two anatomical runs for registration across subjects to standard space: a coplanar T1-weighted FLASH sequence and a high-resolution 3D T1-weighted MPRAGE sequence. An in-plane magnetic field map image was also acquired for EPI undistortion.

### Regions of interest (ROIs)

The hippocampus ROIs were calculated for each participant in subject space using automatic segmentation in Freesurfer [38]. ROIs of hippocampus subfields CA1 and CA2/3/DG were defined from a probabilistic atlas of the medial temporal lobe [39], projected into subject space for analyses.

### fMRI preprocessing

Functional runs were preprocessed using FEAT (FMRI Expert Analysis Tool) Version 5.98, part of FSL (FMRIB’s Software Library, www.fmrib.ox.ac.uk/fsl), including: removal of first four volumes, motion correction using MCFLIRT (individual runs with excessive estimated motion were excluded); fieldmap-based EPI unwarping using PRELUDE+FUGUE; slice-timing correction using Fourier-space time-series phase-shifting; non-brain removal using BET; spatial smoothing using a Gaussian kernel of FWHM 5mm; and high-pass temporal filtering using a 64s-sigma Gaussian kernel. Functional runs were registered with FLIRT to the FLASH image, the MPRAGE image, and an MNI standard brain with interpolation to 2mm isotropic voxels.

### General Linear Model (GLM) for calculation of satellite templates

We modeled the evoked response to individual satellites across a dataset concatenating the 8 preprocessed runs for satellite-viewing periods. The GLM was fit using FILM with local autocorrelation correction in FSL. The model contained a regressor for each of the 15 satellites. Each regressor had a delta function at every presentation of the satellite, excluding repetitions added for the one-back task, convolved with a double-gamma hemodynamic response function. There were also 8 regressors indicating which run the data at that TR corresponded to. The resulting parameter estimates reflected the response of all voxels to each individual satellite, which we call the satellite’s *template*.

### Replay analysis

Within each session for each participant, we compared each of these templates to the pattern of activity at each TR of the rest period to find potential replay events. After preprocessing the data from the rest period as described above, the time series was low-pass filtered through a convolution with a three-point-width Hamming window. This transformation makes the data frequency closer to that of the hemodynamic response function, serving as a replacement for event-related convolution in this scenario where event timing is unknown. We next converted to percent signal change and excluded any voxels with variance more than three SDs from the mean. Then, for all voxels within an ROI or a searchlight (defined as described below), we computed the Pearson correlation between each template and the pattern of activity at each TR of the rest period (Fig. 2a). This resulted in a matrix of positive and negative values for 15 satellites by 191 TRs in the rest period (Fig. 2b, left).

**Figure 2.**
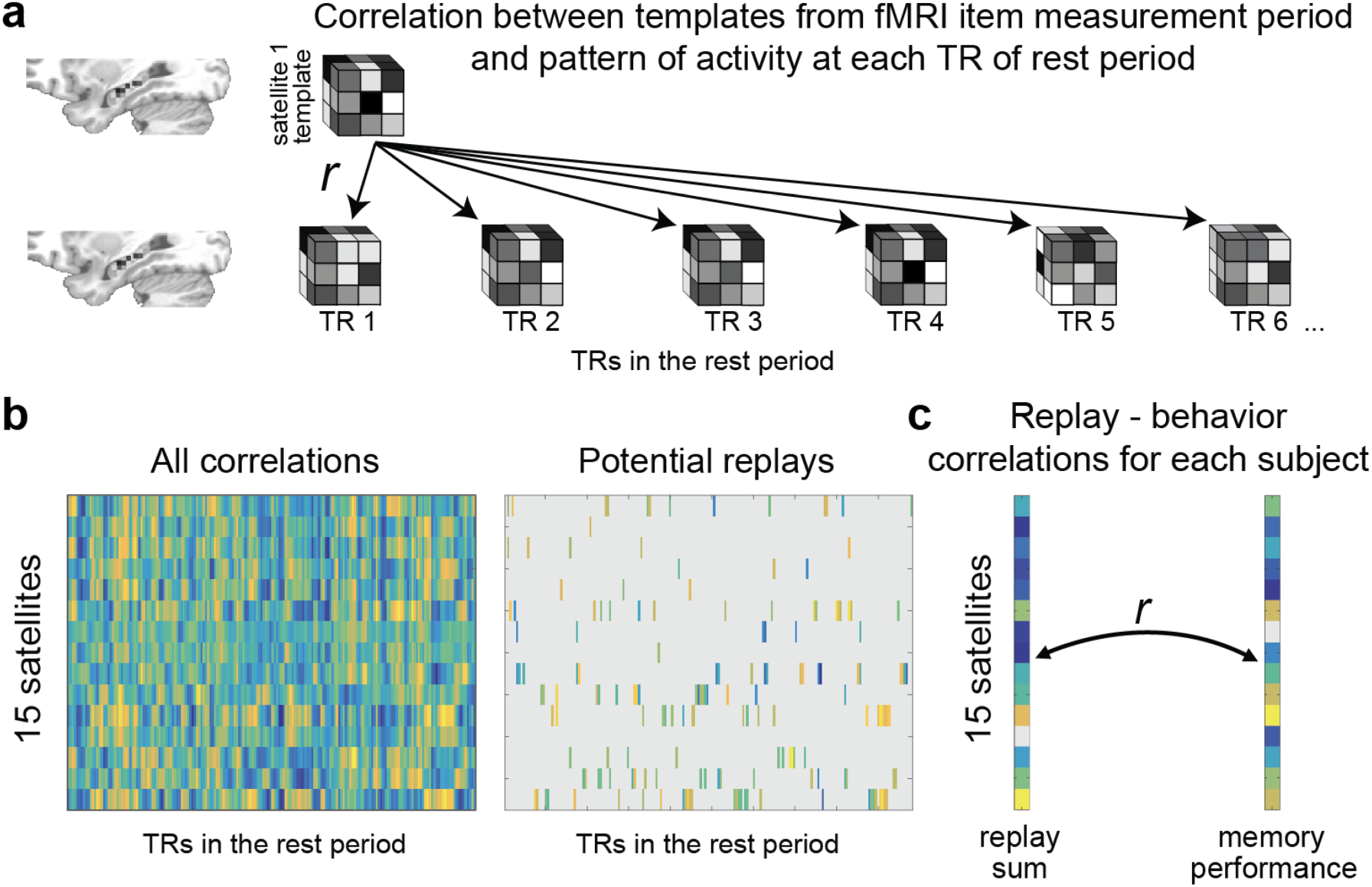
Methods. (**a**) Each satellite’s template (satellite 1 shown as example) consists of a pattern of beta weights across all voxels in the hippocampus. The template is correlated with the preprocessed pattern of activity across all voxels in the hippocampus at each TR of the rest period. (**b**) This results in a matrix of correlation value between all templates and all rest period TRs, which is thresholded to reflect “potential replays.” (**c**) The potential replay activity for a given template is summed across TRs and then correlated with memory performance.

To narrow this matrix down to potential replay events, we only considered activity with strong correspondence between templates and rest period activity. At each TR, we defined strong correspondence as more than 1.5 SDs above the mean across the 15 satellites (1.5 value based on [7]). We chose this approach to increase our chances of seeing satellite-specific replay, as opposed to a brain state at a given TR that resembles satellites in general, while still allowing for the possibility that multiple satellites (or none) are replayed at a given TR.

After thresholding the matrix at each TR (Fig. 2b, right), we summed the total amount of potential replay activity across all TRs for each satellite, and then correlated these values with that individual’s memory for the satellites (Fig. 2C). Across-subject statistics were calculated on these Fisher-transformed correlation values.

### Representation of category structure

Within the hippocampus and subfield ROIs, we calculated all pairwise correlations between templates corresponding to satellites from the same category and all correlations between templates for satellites from different categories, and subtracted the average between-category correlation from the average within-category correlation. This provides an estimate of the extent to which objects from the same category are represented similarly.

### Searchlight analyses

We additionally ran the replay and representation-of-category-structure analyses within every 3 x 3 x 3 voxel cube in the brain. We assigned the final replay-behavior correlation value (Fig. 2c) for the replay analysis, or the correlation difference value for the category-structure analysis, to the center voxel of each searchlight, and then projected these maps to MNI space to allow comparison across participants. We used the *randomise* function in FSL to perform permutation tests to test for reliable clusters, using a cluster formation threshold corresponding to *p*=0.01.

### GLM for univariate change effects

We also ran a GLM to test for overall univariate differences in activity level between the first and second session. Instead of one regressor for each satellite, the model had one regressor with a delta function at every image presentation. We subtracted the map for Session 1 from the map for Session 2, and assessed cluster reliability for Sleep and Wake groups separately as well as contrasting Sleep and Wake groups.

## Results

### Sleepiness survey

KSS scores did not differ across Sleep and Wake conditions for the beginning of Session 1 (mean Sleep = 4.042; mean Wake = 4.417; *t*[22]=0.462, *p*=0.649), end of Session 1 (mean Sleep = 6.167, mean Wake = 5.333, *t*[22]=1.299, *p*=0.207), beginning of Session 2 (mean Sleep = 4.083, mean Wake = 4.083, *t*[22]=0, *p*=1), or end Session 2 (mean Sleep = 4.250; mean Wake = 5.208, *t*[22]=1.101, *p*=0.283), suggesting that there were no alertness differences between groups due to time of day.

### Training and one-back performance

Participants trained for an average of 121.5 trials (SD=83.2), including repetition trials for incorrect choices. Average proportion correct on the last training block was 0.747 (SD=0.088). Detection performance on the one-back cover task in the scanner was excellent (mean A' = 0.916, SD = 0.060).

### Test performance

Performance on the first test was not different for subjects in the Sleep vs. Wake groups in unique, shared, or novel item features (*p*s>0.564). Performance was also not different within each group for unique and shared features, indicating a successful match based on the different feature-type query frequencies used in training (mean Sleep unique = 0.701, mean Sleep shared = 0.677, *t*[11]=0.444, *p*=0.666; mean Wake unique = 0.695, mean Wake shared = 0.715, *t*[11]=0.686, *p*=0.507).

Based on our prior work using this paradigm [32], we predicted that the Sleep group would improve in memory for shared features from the first to second session, the Wake group would decline in memory for shared and unique features, and the Sleep group would improve more than the Wake group for shared and unique features. We thus employed one-tailed t-tests in these analyses. As expected, we found a reliable difference between Sleep and Wake groups in improvement from the first to second session in overall memory (Fig. 3; *t*[22]=2.110, *p*=0.023, one-tailed), and a reliable difference between groups for unique visual (*t*[22]=2.190, *p*=0.020, one-tailed) and shared visual features (*t*[22]=1.995, *p*=0.029, one-tailed).

**Figure 3.**
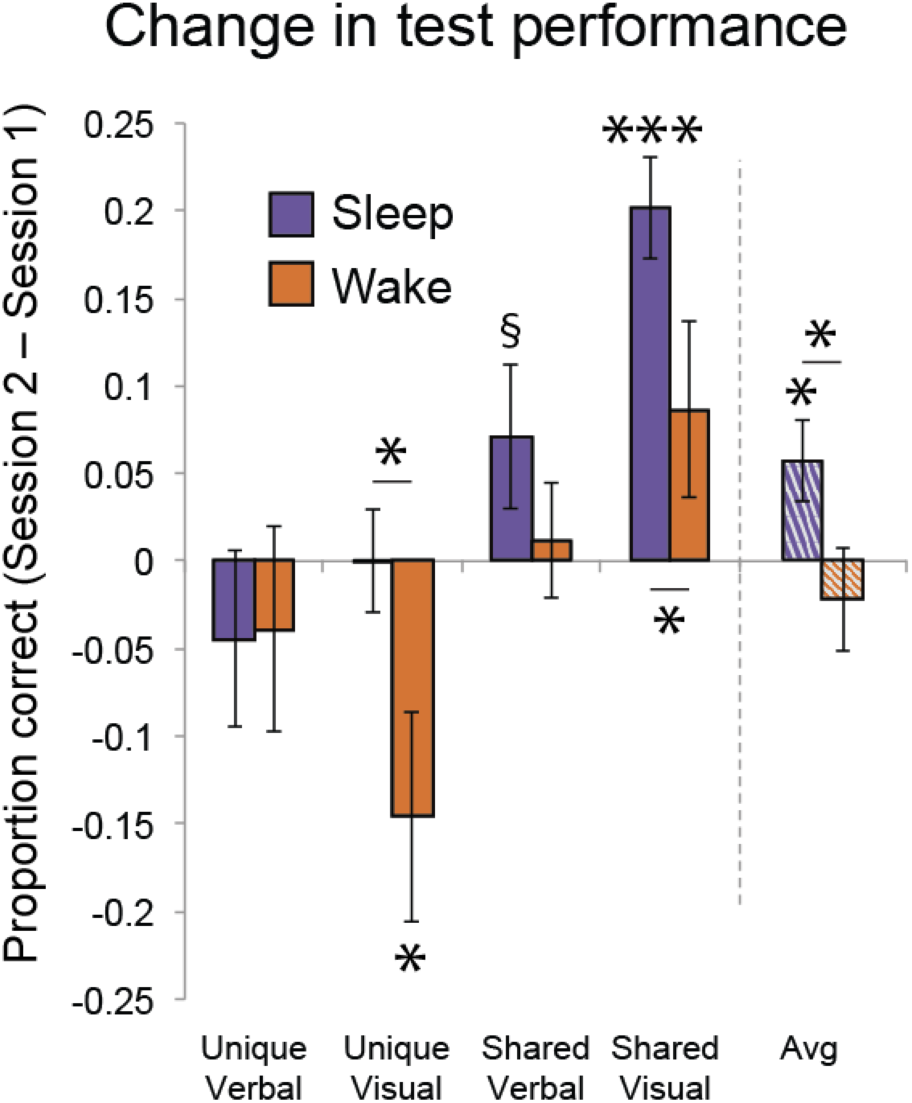
Behavioral results. Change in proportion correct from first to second session for Sleep and Wake group, for different feature types and the average across feature types (Avg). § *p*<0.1, * *p<*0.05, *** *p*<0.001. Asterisks above horizontal lines show significant differences between conditions; asterisks without bars indicate where conditions differ from zero. Error bars denote ± 1 SEM.

The Sleep group showed significant improvement in overall memory (collapsing across shared and unique features) from Session 1 to Session 2 tests (Fig. 3; mean improvement in proportion correct = 0.057; *t*[11]=2.498, *p*=0.015, one-tailed), while the Wake group showed a nonsignificant decrement in performance (mean = −0.022, *t*[11]=0.731, *p*=0.240, one-tailed). As in our prior study, memory for shared features in the Sleep group increased reliably, while memory for unique features stayed constant (mean change for shared = 0.137, *t*[11]=4.187, *p*=0.001, one-tailed; mean change for unique = −0.022, *t*[11]=0.693, *p*=0.503). In the Wake group, there was a nonsignificant improvement in memory for shared features (mean = 0.049, *t*[11]=1.333, *p*=0.896, one-tailed), and a decrease in memory for unique features (mean = −0.092, *t*[11]=-2.081, *p*=0.031, one-tailed). Note that participants were exposed to the visual features of the satellites in the scanner between the two behavioral tests, which could have affected absolute levels of change from Session 1 to Session 2 within each group; however, this issue does not affect Sleep/Wake comparisons, as visual feature exposure was matched across these conditions.

Change in novel item feature performance was similar for the two groups, as we had found in our prior study. There was a marginal increase in performance in both groups (Sleep mean = 0.120, *t*[11]=1.995, *p*=0.071; Wake mean = 0.074, *t*[11]=2.000, *p*=0.071). Since we did not measure the representation of novel items in the scanner, we do not consider these items in further analyses.

### Correlation between memory performance and subsequent hippocampal replay within Session 1

To investigate the relationship between Session 1 performance and subsequent hippocampal replay, we correlated, for each subject, memory for each satellite in Session 1 with the sum of (putative) hippocampal replay activity for that satellite during Session 1 (see Fig. 2 for methods). We then assessed the mean value of these correlations across subjects. We collapsed across Sleep and Wake groups for these analyses. Collapsing across left and right hippocampus, we found reliably negative correlations (mean=-0.145, *t*[23]=3.442, *p*=0.002), which were significant in both left (mean=-0.113, *t*[23]=2.255, *p*=0.034) and right (mean=-0.177, *t*[23]=2.664, *p*=0.014) hippocampus individually, indicating that replay was strongest for the satellites that were remembered worst on the preceding test (Fig. 4).

**Figure 4.**
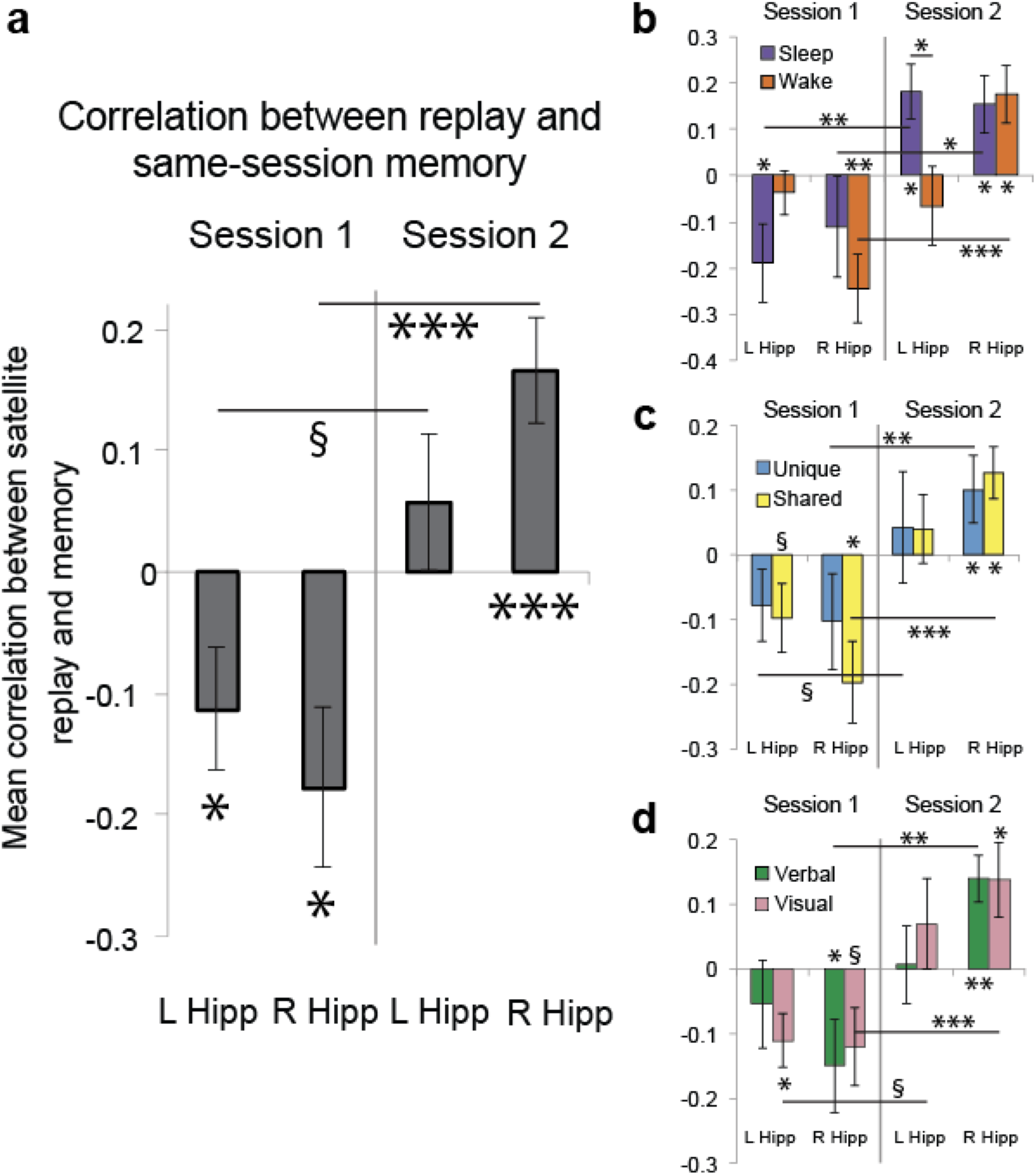
Within-session replay-behavior relationships. (**a**) Correlation between memory in Session 1 and replay in Session 1, and between memory in Session 2 and replay in Session 2, in left and right hippocampus. (**b**) Same data, broken down by Sleep and Wake groups. (**c**) Same data, broken down by memory for unique and shared features. (**d**) Same data, broken down by memory for verbal and visual features. § *p*<0.1, * *p<*0.05, ** *p<*0.01, *** *p*<0.001. Error bars denote ± 1 SEM.

### Correlation between hippocampal replay and subsequent memory performance within Session 2

We did the same analysis as above, now correlating Session 2 replay with subsequent Session 2 memory. Here we found positive correlations that were reliable when combining correlations across the left and right hippocampi (mean=0.112; *t*[23]=3.262, *p*=0.003) and also in the right hippocampus on its own (mean=0.165, *t*[23]=3.841, *p*=0.0008; mean left hippocampus = 0.058, *t*[23]=1.021, *p*=0.318). This indicates that satellites that are replayed more in Session 2 are subsequently better remembered. The correlations were more positive in Session 2 than Session 1 (combining correlations across left and right hippocampi: *t*[23]=4.785, *p*=0.00008; left: *t*[23]=2.039, *p*=0.053; right: *t*[23]=5.164, *p*=0.00003).

### Effects of group, hemisphere, and feature type

To assess whether these effects differed by Sleep vs. Wake group, left vs. right hippocampus, verbal vs. visual information, and unique vs. shared features, we tested within each session whether the results were different for each variable collapsing across the other variables. We opted for this approach because breaking the results down by combinations of these features leads to cells with correlations that cannot be computed, in cases where subjects performed at floor or ceiling on all items within that item type (though there are still a few cases where subjects performed perfectly and had to be excluded from a particular analysis, as reflected in the variation in *df* below). We found no effects of any of these variables. In both sessions, left was not different than right hippocampus (Session 1: *t*[23]=0.773, *p*=0.447; Session 2: *t*[23]=1.453, *p*=0.160), the Sleep group was not different than Wake (Session 1: *t*[22]=0.368, *p*=0.717; Session 2: *t*[22]=0.764, *p*=0.453), verbal was not different than visual (Session 1: *t*[22]=1.001, *p*=0.328; Session 2: *t*[21]=0.756, *p*=0.458), and shared was not different than unique (Session 1: *t*[23]=0.203, *p*=0.841; Session 2: *t*[20]=0.693, *p*=0.496).

### Cross-session replay-behavior relationships

Session 1 replay was correlated with Session 2 behavior (mean=-0.136, *p*=0.039, collapsed across right and left hippocampus; correlation between Session 2 replay and Session 1 behavior mean=0.077, *p*=0.135), which is likely due to the fact that behavior is correlated across sessions (mean correlation=0.394, *p*<0.0001). This relationship did not withstand regressing out within-session behavior (mean coefficient=-0.027, *p*=0.175; for Session 2 replay, mean coefficient=-0.651, *p*=0.964), whereas within-session behavior-replay relationships largely remained when regressing out other-session behavior (mean coefficient for Session 1 behavior predicting Session 1 replay after regressing out Session 2 behavior=-1.22, *p*=0.066; mean coefficient for Session 2 behavior predicting Session 2 replay after regressing out Session 1 behavior: 1.339, *p*=0.051). Memory and replay are thus most closely related within the same session.

### Cross-session replay-replay relationships

For each individual, we calculated the correlation between amount of replay of each of the 15 satellites in Session 1 and amount of replay of each of the 15 satellites in Session 2. These correlations were not reliably different from zero across subjects (*t*[23]=0.871, *p*=0.393, collapsed across right and left hippocampus).

### Correlation between hippocampal replay and memory performance change from Session 1 to Session 2

We next separated Sleep and Wake groups and asked whether replay in either session relates differently to behavioral change over time for subjects who slept vs. did not sleep between sessions. Because prior work strongly predicts that replay should improve performance over sleep, we again employed one-tailed t-tests in the analysis of the Sleep group, and in the comparison between Sleep and Wake groups. We correlated replay for each satellite in each session with change in performance for that satellite (Session 2 – Session 1 performance). We found that, across hemispheres of the hippocampus and across sessions, replay in the Sleep group was indeed positively related to memory improvement (Fig. 5; mean=0.093, *t*[11]=2.146, *p*=0.028, one-tailed), and that this effect was reliably greater than in the Wake group (*t*[22]=2.522, *p*=0.010, one-tailed), which had a numerically negative relationship (mean=-0.066, *t*[11]=1.441, *p*=0.177). For particular sessions and hemispheres, there was a marginal negative effect in left hippocampus in Session 1 in the Wake group (mean=-0.134, *t*[11]=1.814, *p*=0.097), which was significantly lower than in the Sleep group (*t*[22]=1.782, *p*=0.045, one-tailed), and a positive effect in the Sleep group in the right hippocampus in Session 2 (mean=0.186, *t*[11]=2.289, *p*=0.022, one-tailed).

**Figure 5.**
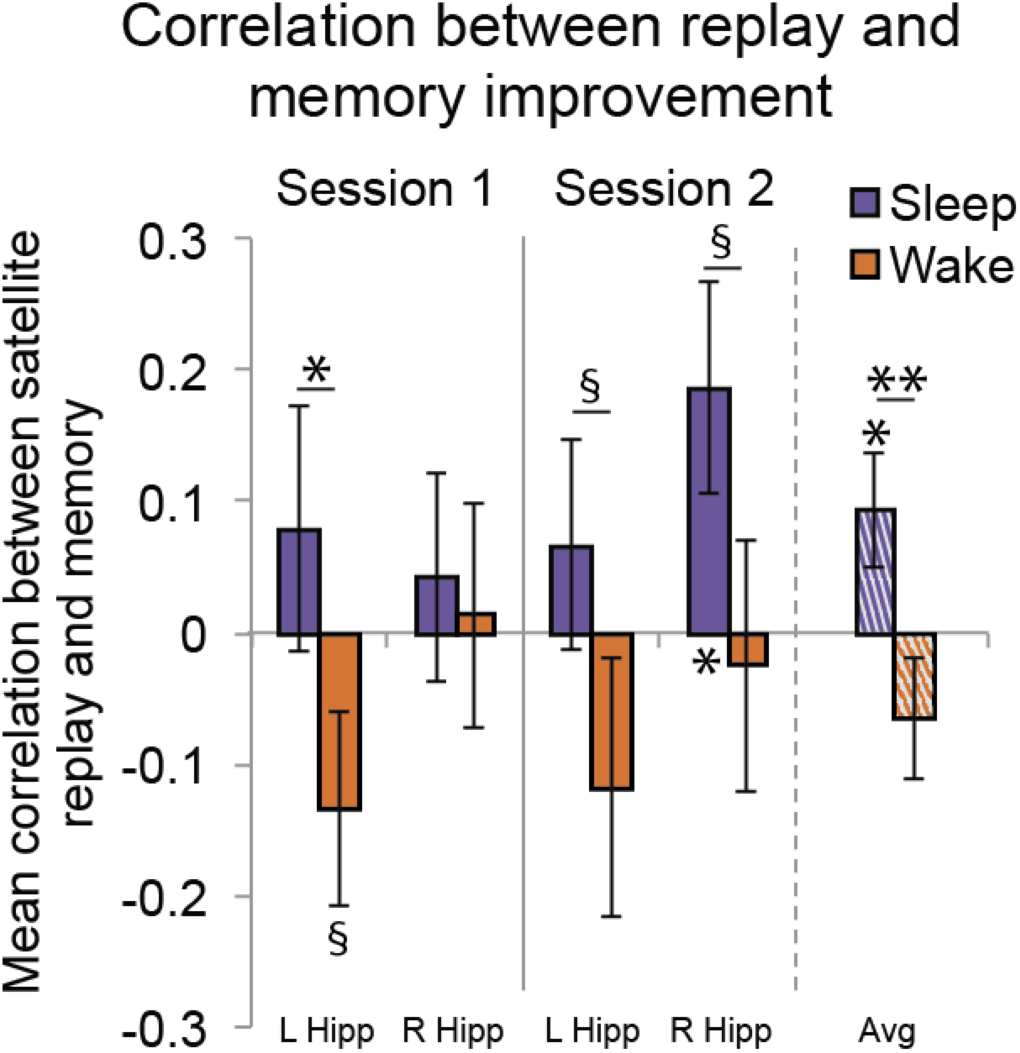
Across-session replay-behavior relationships. Correlation between replay in each session and improvement in memory from the first to second session. § *p*<0.1, * *p<*0.05 ** *p*<0.01. Error bars denote ± 1 SEM.

### Overall replay

Our design did not allow us to assess true overall replay amounts, as we did not have a baseline scan session prior to participants learning about the satellites. However, we can compare overall replay amounts across sessions or groups. Overall replay amount did not differ between Session 1 and 2, collapsed across groups (both hippocampi: *t*[23]=0.206, *p*=0.839; left: *t*[23]=0.797, *p*=0.434, right: *t*[23]=0.324, *p=*0.749). Overall replay amount also did not differ between Sleep and Wake groups within Session 1 (both hippocampi: *t*[22]=1.351, *p*=0.191; left: *t*[22]=0.654, *p*=0.520; right: *t*[22]=1.454, *p=*0.160) or Session 2 (both hippocampi: *t*[22]=0.633, *p*=0.534; left: *t*[22]=0.503, *p*=0.620; right: *t*[22]=1.390, *p=*0.178).

### Representation of category structure

To test whether the hippocampus was sensitive to the category structure of the stimuli, we assessed whether satellites from the same category were represented more similarly than satellites from different categories, collapsing across groups and sessions. The hippocampus overall did not show any sensitivity to category structure (both hippocampi: *t*[23]=0.341, *p*=0.737; left: *t*[23]=0.845, *p*=0.407; right: *t*[23]=1.600, *p*=0.123). Considering separate subfields, however, a reliable effect was observed in CA1 (both hemispheres: *t*[23]=2.336, *p*=0.029; left: *t*[23]=1.685, *p*=0.106; right: *t*[23]=2.345, *p*=0.028) but not CA2/3/DG (both hemispheres: *t*[23]=0.612, *p*=0.547; left: *t*[23]=1.068, *p*=0.297; right: *t*[23]=0.359, *p*=0.723), consistent with a previous proposal about the role of CA1 in representing structured information [40]. None of the results differed by session or group (*p*s>0.196).

### Wholebrain analyses

We ran exploratory wholebrain searchlights looking for other regions where item replay during Session 1 or Session 2 might relate to behavior within that session, or regions where replay during Session 1 or Session 2 might relate to the change in performance from Session 1 to 2. No regions survived multiple comparisons correction.

We also ran a searchlight to test whether areas outside the hippocampus might represent the category structure (as found in hippocampal subfield CA1). We found two significant clusters, both associated with visual processing, identified according to a probabilistic atlas of the visual system [41]: One in visual cortex (corrected *p*=0.0002), spanning V1-V4, and another in frontal cortex (corrected *p*=0.031), partially overlapping with the frontal eye fields and falling within regions of the inferior frontal sulcus involved in spatial vision [42]. Responses in these areas likely reflect the fact that objects in the same category share visual features. There were no differences in category structure between sessions or groups in this wholebrain analysis.

We next tested whether there were any changes in overall, univariate activity level from the first to second session. While there were no effects that survived cluster correction for the Wake group, or for Sleep contrasted with Wake, we did find a reliable cluster in the medial prefrontal cortex (mPFC) that increased in activity from Session 1 to 2 in the Sleep group (corrected *p=*0.030; coordinates of center of gravity: 45.2, 87.8, 35.8; extent=521 voxels).

### Assessing the role of attention

There is a potential alternative explanation for the negative correlation between memory and subsequent replay in Session 1 due to the presence of the intervening fMRI measurement period: If participants attend more during the measurement period to the stimuli that they had just performed poorly on, this could cause persistence of the representations of those attended satellites into the subsequent rest period. To test this idea, we assessed whether satellites with worse average memory performance in Session 1 had higher BOLD activity in a canonical attention network (“attention” network from neurosynth.org using reverse inference: *attention_pFgA_z_FDR_0.01.nii*) during the subsequent measurement period. We found no evidence for a relationship (mean correlation = 0.014, *t*[23]=0.253, *p*=0.803). We also tested for this effect in the hippocampus and found no relationship there (mean correlation = −0.003; *t*[23]=0.060, *p*=0.952). Next, we ran the analysis at every voxel in the brain and assessed whether any areas showed reliable cluster-corrected results. There were no reliable clusters looking across the whole brain, nor with small volume correction within the attention network (all *p*s>0.686). We therefore conclude that attention to poorly-remembered stimuli during the measurement period is unlikely to be driving the negative relationship between memory and replay in Session 1.

## Discussion

How does hippocampal replay during quiet rest relate to previous learning and to subsequent memory? To address these questions, we taught participants the features of 15 objects and assessed replay of individual object representations before and after memory tests. We did extensive repeated exposure of the objects during fMRI scanning to maximize our ability to create a high-fidelity template that could be used to track replay. We found that objects that were initially remembered poorly were subsequently replayed more in the hippocampus, while in a second session, objects that were replayed more were subsequently better remembered. Furthermore, replay was related to change in memory from the first to second test only for participants that slept between the two sessions. These results address the four questions raised in the introduction, with important implications for our understanding of consolidation of object knowledge.

### Individual items are replayed in the human hippocampus

While prior work has shown item-specific replay outside the hippocampus [7-9] and coarser replay within the hippocampus [10, 14, 46], the current results provide the first evidence for item-specific replay in the human hippocampus. This demonstration of item-specific replay is grounded in the observed correlations (computed across-items, within-subjects) between replay and memory performance.

### Replay prioritizes weaker memories

Prior fMRI studies showing a positive relationship between replay and subsequent memory [7-16] fit with the idea that replay causes better memory. However, these results can also be explained in terms of stronger memories being replayed more (without there necessarily being a causal relationship between replay and subsequent memory). Our results from the second session fit with this latter story, but the results from the first session do not. Instead, the negative correlation between replay and preceding memory in the first session suggests the use of a prioritization algorithm that focuses on weaker memories. This is consistent with prior findings from this paradigm [32], where weaker items benefitted from a short period of sleep (whereas overnight sleep benefitted all items). Taken together, these findings suggest that the brain’s first priority is to process weaker information. A recent study in rodents found that replay during rest was important for gradually-strengthened but not quickly-formed memories [6], perhaps reflecting a similar process. Prioritization of weaker information is also consistent with the finding that encoding difficulty predicts the occurrence of more sleep spindles (which are associated with replay) during a subsequent nap [43] and the general tendency for sleep to benefit information that is more weakly encoded [22-29, cf. 30, 31].

One human fMRI study found that activity patterns in the hippocampus that were strongest during encoding (explaining the most variance in a Principal Component Analysis) persisted into rest and correlated with subsequent memory [10]. This may seem to contrast with our study and others, but these strong patterns did not necessarily correspond to strongly-encoded items—they could in principle relate to a monitoring process associated with weaker memories. Of course, memory strength is not the only dimension that can drive prioritized wake replay; other known dimensions include fear [17] and reward [14, 15, 44].

### Replay is associated with better subsequent memory

In Session 2, items replayed more showed better subsequent memory. This result is consistent with accumulating findings from both the rodent [2, 5] and human literature [7-17, 45] suggesting a functional role for hippocampal replay (i.e., improved consolidation), although—for reasons discussed above—this correlation unto itself is not evidence that replay causes better subsequent memory.

### The relationship of wake replay to processing during sleep

Wake replay in both sessions was associated with greater memory improvement for participants who slept than for those who remained awake between sessions. Thus while wake replay may have its own benefits, it may also lay the groundwork for subsequent sleep-dependent consolidation processes. One hypothesis suggested in other work [47, 48] is that processing during wake rest serves to “tag” memory traces for subsequent replay during sleep. Alternatively, it may be that some other factor occurring prior to rest (during initial encoding) drives prioritization during both wake and sleep replay, and that the larger effects observed in the Sleep group arise because those participants encountered less wake-based interference, or more replay overall during sleep. In either case, some process must occur prior to the rest period that constrains which representations are revisited later. The process cannot simply reflect encoding of an external error signal, since participants did not receive feedback during the test phase in our study. Instead some process sensitive to the strength of the representation generated during test must have provided the signal that subsequently led to increased wake replay.

### Systems consolidation

Consistent with a prior study [32], memory for the satellite objects, and in particular the features shared across members of a category, was better after a night of sleep than a day awake. These results may reflect an increased reliance on cortical areas as a result of systems consolidation [49], as cortical areas employ more overlapping representations than the hippocampus and should therefore facilitate memory for shared features [32, 50]. One cortical region often implicated as a consolidation site is the mPFC [51], and our whole-brain univariate analysis found increased activation across sessions in that region for the Sleep but not the Wake group. We did not find any evidence that the hippocampus was less involved in this task from the first to second session in the Sleep group [cf. 52], though any decreased reliance on the hippocampus may take more time.

### Conclusion

Our findings suggest that wake replay in the hippocampus does not simply reflect the strongest representations rising to the surface, but instead is adaptive, prioritizing memories that most need strengthening. The results provide the first evidence for hippocampal replay of individual memories in humans, and add to a growing literature suggesting that wake replay is associated with improved subsequent memory. The findings also point to a relationship between wake replay and the memory processing that occurs over a night of sleep, a promising dynamic to explore in future research.

## Acknowledgments

We thank Luis Piloto for help with initial analyses and subject running and Michael Arcaro for advice on processing the rest data and identification of visual areas. This work was supported by: NIH NINDS F32-NS093901 (ACS); NSF GRFP (EAM); NIH NIA R01-AG046646 (SCM); NSF BCS-1439210 (SCM); NIH NIMH R01-MH069456 (KAN).

## Footnotes

1 Though the word “replay” is sometimes reserved for the sequential replay of place cell activity observed in rodents, we use the term more broadly to mean neural reactivation of recent experience, including that observed in fMRI.

